# Distinguishing between models of mammalian gene expression: telegraph-like models versus mechanistic models

**DOI:** 10.1101/2021.06.08.447592

**Authors:** Svitlana Braichenko, James Holehouse, Ramon Grima

**Affiliations:** School of Biological Sciences, University of Edinburgh, United Kingdom; School of Informatics, University of Edinburgh, United Kingdom

**Author notes:** These authors contributed equally (Svitlana Braichenko, James Holehouse).

## Abstract

Two-state models (telegraph-like models) have a successful history of predicting distributions of cellular and nascent mRNA numbers that can well fit experimental data. These models exclude key rate limiting steps, and hence it is unclear why they are able to accurately predict the number distributions. To answer this question, here we compare these models to a novel stochastic mechanistic model of transcription in mammalian cells that presents a unified description of transcriptional factor, polymerase and mature mRNA dynamics. We show that there is a large region of parameter space where the first, second and third moments of the distributions of the waiting times between two consecutively produced transcripts (nascent or mature) of two-state and mechanistic models exactly match. In this region, (i) one can uniquely express the two-state model parameters in terms of those of the mechanistic model, (ii) the models are practically indistinguishable by comparison of their transcript numbers distributions, and (iii) they are distinguishable from the shape of their waiting time distributions. Our results clarify the relationship between different gene expression models and identify a means to select between them from experimental data.

## 1 Introduction

One of the most popular models of gene expression is the *telegraph model*, a two-state model where genes are assumed to be either *on* or *off*, being able to produce mature messenger RNA (mRNA) in the on state and having no mature mRNA production in the off state [1–3]. Since gene expression is inherently stochastic [4], mathematical models of the telegraph model often employ probabilistic modelling techniques such as the chemical master equation [5, 6] or the stochastic simulation algorithm (SSA) [7]. By fitting the steady-state analytical solution of the telegraph model to experimentally measured distributions of the number of cellular mRNA in single cells, several studies have estimated the rates of gene switching and of initiation for several mammalian genes [8–12]. However, mapping cellular mRNA number to the underlying transcription kinetics is difficult because fluctuations in this number reflect noise due to many processes downstream of transcription [13, 14].

In contrast, the number of actively transcribing RNA polymerase II (Pol II) on a gene is not subject to these complex processes, and hence reveals more information on the details of transcription [15–17]. Therefore, unlike mature mRNA statistics, fluctuations of actively transcribing Pol II provide a direct readout of transcription. Since the speed of actively transcribing Pol II is approximately constant along a gene and since its premature detachment is not frequent, it follows that the loss of actively transcribing Pol II (leading to the production of a mature mRNA transcript) cannot be described by a first-order reaction (as assumed in the telegraph model for cellular mRNA). Rather it is much better captured by a delayed degradation reaction where the removal of an actively transcribing Pol II occurs after a fixed elapsed time since its binding to the promoter. A recent paper [14] has modified the telegraph model to account for the aforementioned speciality, a model that we shall refer to as the *delayed telegraph model*. This alternative two-state model, unlike the telegraph model, is non-Markovian; while its mathematical analysis is complex, it can be solved exactly in steady-state to obtain distributions of the number of bound Pol II. Transcriptional parameters can then be obtained by fitting these distributions to those obtained experimentally using electron microscopy [18] or nascent RNA sequencing [19]. Alternatively, since each actively transcribing Pol II has attached to it an incomplete nascent mRNA, one can also use the delay telegraph model to numerically calculate the steady-state distribution of nascent mRNA numbers which can then be fit to distributions obtained using single-molecule fluorescence in situ hybridization (smFISH) [20].

Despite their success in predicting distributions of transcript numbers that match those calculated from experimental data, it is important to remember that both the telegraph model and the delayed telegraph model do not include a description of all the key rate limiting steps. In the past decade, several experimental papers have shown the necessity of including Pol II pausing and release in models of transcription. Bartman et al. [21] show experimentally that it is the release of Pol II from the pausing state, and not the Pol II recruitment rate, that is a key control point for gene expression. In fact, it is found universally amongst all metazoan genes that the rate of release of Pol II from pausing is the rate limiting step in transcription [22]. In mammalian cells the release of Pol II from the paused state is dependent on the activity of several molecules, including the transcription elongation factor P-TEFb [22–24]. Specifically in embryonic stem cells, ChIP-Seq data has revealed that Pol II peaks near genes are at the promoter-proximal region, and that inhibiting the P-TEFb causes Pol II to remain in the promoter-proximal region genome-wide [24]. Figures 1 and 2 in [22] provide a good overview of the key step of transcription, including Pol II pausing and release. The mechanism of Pol II pausing, in addition to the binding of Pol II and other transcription factors to the promoter, provides two layers of control over the production of nascent and mature mRNA. It is also found that expressed genes without a peak of paused Pol II in one cell type can acquire pausing in a different cell type, therefore genes have the potential of being regulated by proximal pausing even when the Pol II pausing peak is absent [23]. Clearly, if Pol II pausing and release is such a key feature of transcriptional models, the current ambiguity of the mechanisms’ roles in the standard and delayed telegraph models is a problem in need of a solution.

**Figure 1:**
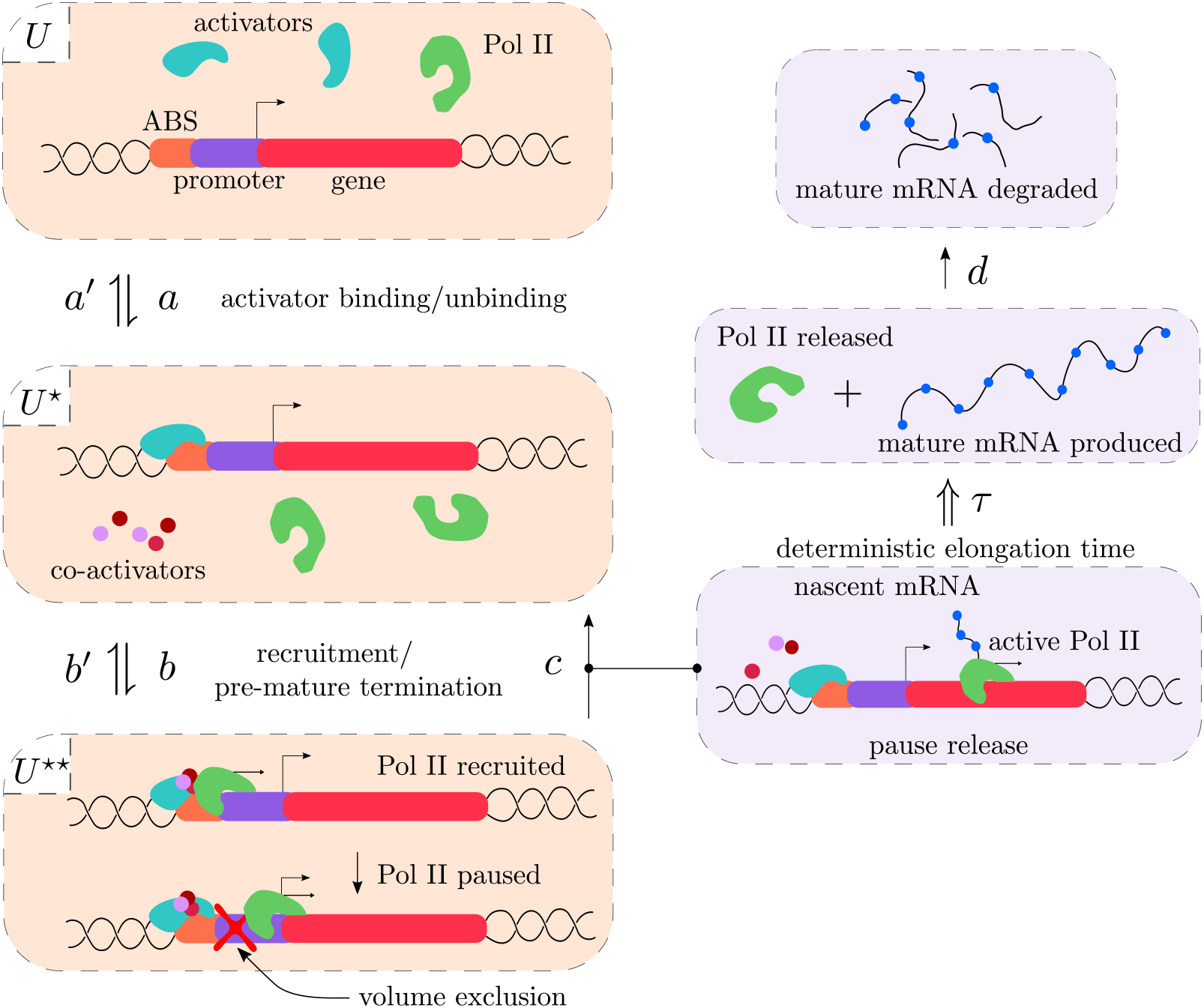
Illustration of system (1). The *U* state describes the state where both the activator binding site (ABS) and the promoter are unbound. In the *U*\* state, the activator is bound to the ABS meaning the Pol II can bind to the promoter. Pol II has been recruited to the promoter and pauses in state *U*\*\*. Transitions from *U*\*\* to *U*\* either result from premature termination or else pause release and the subsequent production of an actively transcribing Pol II. Elongation (and termination) takes a deterministic time *τ* after which the mature mRNA is produced. The latter is subsequently degraded in the cytoplasm. For more details see the main text.

**Figure 2:**
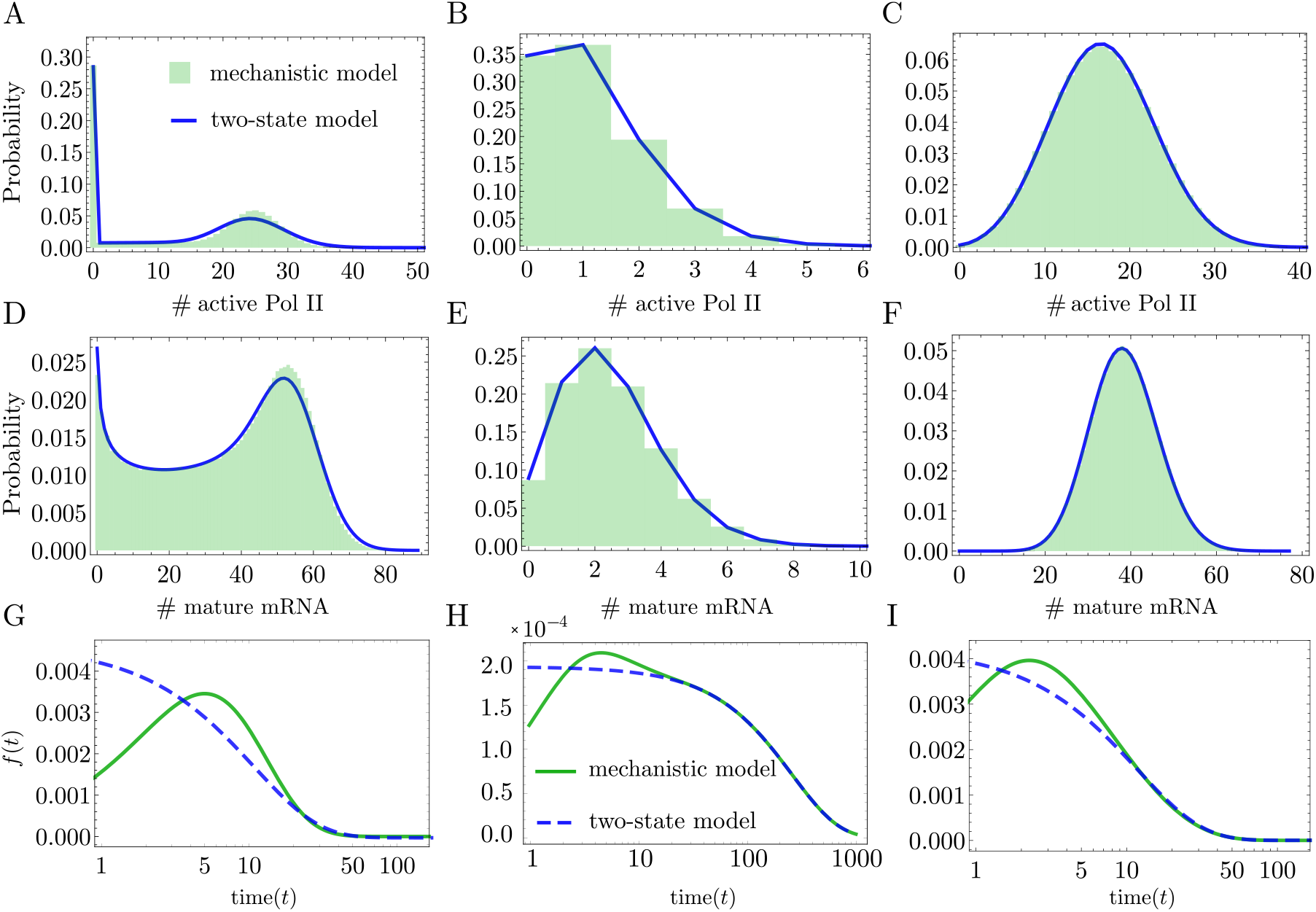
Comparison between the molecule number distributions of active Pol II and mature mRNA distributions of the two-state (reduced) models ((2) and (3)) and the mechanistic model (1). For a particular choice of parameters of the mechanistic model, Eq. (7) gives the two-state model parameters. In (A-C), for three different parameter sets (one per column), we show that the mechanistic model describing the number of active Pol II molecules is well approximated by the exact solution of the delay telegraph model [14] evaluated with the effective parameters. (D-F) show a similar level of agreement between the mechanistic and two-state models but instead for the mature mRNA distributions, where the two-state model is now the telegraph model whose exact solution can be found in [1]. Distributions of the mechanistic model were obtained using the delay SSA (Algorithm 2 of [49]) with 10^4^ samples. In (G-I) we show the corresponding distributions of the waiting time between two consecutive active Pol II (or mature mRNA) production events for the two-state and mechanistic models. The waiting time distributions for the mechanistic and two-state models are calculated by taking the inverse Laplace transform of Eq. (26) and Eq. (39) respectively. Clearly, the models can be distinguished through their waiting time distributions *even when their number distributions are almost indistinguishable*. Parameters of the mechanistic model and the corresponding effective parameters for two-state models are: (A) *a* = 0.001 s^−1^, *a*′ = 0.001 s^−1^, *b* = 0.16 s^−1^, *b*′ = 0.016 s^−1^, *c* = 0.24 s^−1^ mapped to *σ_u_* = 0.0007 s^−1^, *σ_b_* = 0.001 s^−1^, *ρ* = 0.092 s^−1^; (B) *a* = 0.144 s^−1^, *a*′ = 0.032 ^s−2^, *b* = 0.016 s^−1^, *b*′ = 0.56 s^−1^, *c* = 0.24 s^−1^ mapped to *σ_u_* = 3.8 10^−8^ s^−1^, *σ_b_* = 0.002 s^−1^, *ρ* = 0.004 s^−1^; (C) *a* = 0.032 s^−1^, *a*′ = 0.032 s^−1^, *b* = 0.16 s^−1^, *b*′ = 0.016 s^−1^, *c* = 0.32 s^−1^ mapped to *σ_u_* = 0.012 s^−1^, *σ_b_* = 0.029 s^−1^, *ρ* = 0.086 s^−1^. The mature mRNA decay rate is *d* = 0.0016 s ^−1^, and the delay time due to elongation is *τ* = 273.62 s.

Thus far, the modelling literature contains few studies where transcription is modelled incorporating Pol II pausing and release. One model, found in [25], includes pausing and release in a three-state gene model based on the findings of [21], where the three states represent (i) an inactive gene state *D*_0_, (ii) a “burst initiated” state *D*_10_ where the gene is bound to transcription factors and enhancers, and (iii) a gene state *D*_11_ in which the Pol II is bound and paused. Mature mRNA is produced in the transition from *D*_11_ → *D*_10_; this reaction should actually produce nascent mRNA but in this model, it is assumed that the nascent lifetime is so short that a nascent mRNA description can be ignored. By ignoring nascent mRNA fluctuations and assuming that the pausing and unpausing of the Pol II is very fast, it was shown in [25] that the mature mRNA distribution from this model is well approximated by that from the telegraph model. Two other recent studies [26, 27] also explore similar models albeit in the context of transcription reinitiation [28].

In summary, it is currently not so clear why the telegraph model is so successful in fitting experimental mature mRNA distributions, even though it misses important reaction steps which are key control points for gene expression. It is unclear if the assumptions made in [25] are necessary to guarantee that the true mature mRNA distribution is well approximated by the telegraph model; it could well be that these are sufficient but not necessary conditions. Since this study did not derive nascent mRNA statistics, nothing can be inferred about the reasons underlying the success of the delayed telegraph model in fitting experimental nascent mRNA distributions. A related and important question still remains: if the two-state and more detailed mechanistic models of transcription cannot be distinguished from distributions of the number of transcripts, is there another statistic that is useful to distinguish between them? In this study we take a first step at answering these questions.

The paper is divided as follows. In Section 2 we introduce the standard and delayed telegraph models (two-state models), as well as a mechanistic multi-state gene model that provides a stochastic description of transcription factor, Pol II and mature mRNA dynamics. Then, in Section 3 we explore the relationship between the two-state and mechanistic models by comparing the distributions of their waiting times between two consecutive transcripts. We show that two-state models can always be told apart from the mechanistic model from the shape of the waiting time distribution, even when they are indistinguishable from a comparison of their number distributions. We also derive conditions under which the moments of the waiting time distribution (up to third order) of the mechanistic model agree with those of two-state models, leading to expressions relating the parameters of the latter with those of the former. In Section 4 we perform a sensitivity analysis using the aforementioned expressions to understand which parameters of the mechanistic model are the parameters of two-state model most sensitive to. This uncovers non-trivial correlations between the parameters of two-state models. In Section 5 we show that the conclusions previously based on waiting time distributions agree with those obtained using model reduction methods based on number statistics. Finally, in Section 6 we conclude the study and discuss our results in the context of the literature.

## 2 Models of transcription

In this section, we start by introducing an effective reaction scheme for a mechanistic model of transcription describing activator, Pol II, and mature mRNA dynamics. Then, we introduce the standard and delayed telegraph models as the two-state models whose dynamics we will attempt to match to that of mature mRNA and actively transcribing Pol II in the mechanistic model, respectively.

### 2.1 A non-Markovian mechanistic model of transcription

The mechanistic model of transcription in metazoan cells that we henceforth consider is defined in terms of the following effective reactions:

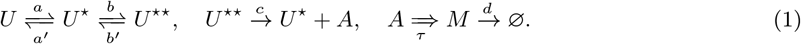

State *U* describes a gene state in which Pol II cannot access the promoter region at the beginning of a gene since activator binding is impaired by chromatin [29, 30]. In contrast, state *U*^⋆^ describes a state where activator binding has reorganised the local nucleosome structure [30], allowing Pol II to access the promoter region along with all transcription factors, co-activators, unphosphorylated Pol II and initiation factors needed for transcription initiation to start. This state is coincident with the dynamic promoter condensate (or transcription factories) proposed in various papers [31–33]. Transcription factors recruit cofactors and Pol II, and hence drive the (reversible) change of state from *U* to *U*^⋆^.

Initiation starts with the binding of Pol II to the promoter; it then pauses promoter-proximally [34]. These processes are modelled by the change of state from *U*^⋆^ to *U*^⋆⋆^, where the latter is the paused state. Once the pause is released, Pol II begins moving away from the promoter region, thus starting productive elongation that leads to a Pol II molecule with a nascent mRNA tail (even paused Pol II has a tail but it is very short and we will hence ignore it). Note that the nascent transcript is not a fully formed mRNA transcript since the length of the tail attached to Pol II increases as elongation progresses. The number of Pol II bound to the gene is equal to the number of nascent mRNA irrespective of their lengths. We call any Pol II undergoing productive elongation as an active Pol II (*A*), which implies that the change of state from paused *U*^⋆⋆^ to unpaused *U*^⋆^ must simultaneously lead to the production of an *A* particle. Note that the binding of another Pol II to the promoter is not possible when there is already a Pol II paused promoter-proximally due to volume exclusion imposed by the latter [35].

Elongation (and termination) finishes after a fixed elapsed time *τ* leading to the detachment of Pol II from the gene and the dissociation of the mRNA tail from Pol II. We hence call the fully formed mRNA a mature transcript *M* and elongation is described by the effective reaction *A ⇒ M* (the double horizontal arrow is here used to denote delayed degradation which occurs after a fixed time *τ*). Note that the change from *A* to *M* cannot be modelled by a first-order reaction because elongation involves the movement of Pol II along the gene with an approximately constant velocity and hence the lifetime of an active Pol II molecule is not exponentially distributed [14, 36].

Note that paused Pol II instead of leading to productive elongation can also undergo premature termination [22] i.e., the paused Pol II releases the short nascent mRNA tail attached to it (which is rapidly degraded) and the polymerase is recycled into the free Pol II pool. These reactions may happen quite often [37, 38]; it is thus quite unlikely that they simultaneously lead to a dissociation of the dynamic promoter condensate since otherwise the efficiency of gene expression would become extremely low. Hence we assume that premature termination leads to a change from the paused state *U*^⋆⋆^ to the unpaused state *U*^⋆^ but do not consider transitions of the type *U*^⋆⋆^ to the non-permissive/inactive state *U*.

Finally, the mature transcripts are removed via various mRNA decay pathways in the cytoplasm [39, 40]. Since many mammalian genes follow single-exponential decay kinetics [41], we model mature mRNA turnover via a first-order reaction of the form *M* → ⌀.

We emphasize that a speciality of our model is the reaction *U*^⋆⋆^ → *U*^⋆^ + *A*. This involves a change of gene state each time a transcription event occurs, whereas common models of gene expression in the literature do not have such a coupling [1, 3, 14, 42–44]. As explained above, the change of state is necessary to model the fine-scale details of the molecular biology, namely the fact that unpausing a Pol II frees the promoter and enables the binding of a new Pol II to it. Unpausing of Pol II is a key rate limiting step since the mapping of Pol II using chromatin immunoprecipitation (ChIP) revealed peaks of Pol II near many promoters [24, 45, 46]. In fact, this accumulation of Pol II near the promoters indicates that the relative rates of premature termination (*b*′) and pause release (*c*) are much slower than rates of recruitment and entry into the paused state (*b*). Since regulatory processes often target rate-limiting steps, the release of paused Pol II has emerged as a central point of gene expression control [21, 22].

A master equation can be written down which describes the stochastic dynamics of this model; its form is quite different than the standard chemical master equation that is popular in the literature of gene expression [6]. The right hand side of the latter equation is only a function of the present time *t*. In contrast, the master equation describing our model has a right-hand side that is a function of not only the present time *t* but of the history of the process in the interval [*t − τ, t*]. This is because of the fixed time *τ* between the release from the paused state and the production of a mature transcript. The dependence of the dynamics of the system on its history means that our model is non-Markovian [47].

### 2.2 Two-state models of transcription: telegraph and delay telegraph models

In the literature, two models are commonly used to (separately) describe active Pol II and mature mRNA dynamics:

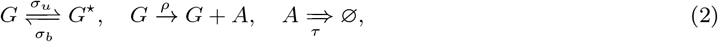

and

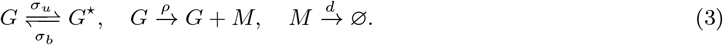

The chemical master equation describing the stochastic dynamics of the systems defined by schemes (2) and (3) were exactly solved in steady-state by Xu et al. [14] and Peccoud and Ycart [1], respectively. Model (3) has also has a transient solution which is reported in [2]. Model (3) is often called the telegraph model of gene expression; by analogy, we shall refer to Model (2) as the delayed telegraph model. Note that the former is a Markov model while the latter is non-Markov in character for the same reason as described above for the mechanistic model.

Clearly, the difference between the two models is how the transcripts are removed from the system: active Pol II is removed after a fixed elapsed time *τ* (due to the termination of elongation which results in a mature transcript), whereas mature mRNA degradation follows first-order kinetics. Both models postulate that at any point in time, the gene is in one of two states: an active state *G* from which transcription can occur and an inactive state *G*\*. As argued by Bartman et al. [21] it is unclear what is the precise biological meaning that should be associated with these two states because the reaction in these models cannot be clearly associated with polymerase processes that are central to transcription.

As we argued in the Introduction, both telegraph and delayed telegraph models have been shown to accurately replicate experimental distributions of mature and nascent mRNA numbers. This leads us to the following question: could it be possible that the stochastic dynamics of our mechanistic model defined by (1) are accurately approximated by these simpler models? To be more precise, is there a set of effective transcriptional parameters *ρ, σ_u_, σ_b_* of the two-state models such that they predict the same (or very similar) distributions of active Pol II and mature mRNA numbers in the mechanistic model. If there is such a set of effective parameters, ideally we would also want expressions showing their relationship to the parameters *a, a, b, b, c* of the mechanistic model.

## 3 Relationship between the parameters of the two-state and mechanistic models

### 3.1 When can the two-state and mechanistic models be matched? A waiting time distribution perspective

In Appendix A and B, we calculate the distribution of the time between the production of two consecutive *M* (*A*) molecules for the mechanistic and (delayed) telegraph models. Using these distributions we can compute the square of the coefficient of variation of the time between two consecutive *M* (or *A*) production events. Throughout the paper, we will refer to this time between production events as the waiting time. In line with previous usage in the single enzyme molecule literature [48], we shall refer to the coefficient of variation of the waiting time distribution as the *randomness parameter*, which is found to be given by:

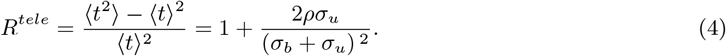

for the *telegraph or delayed telegraph models* and by:

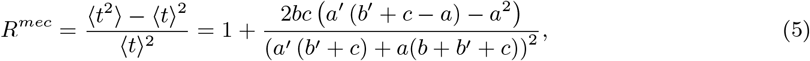

for the mechanistic model. Note that the waiting time statistics for *A* and *M* in the two-state models are the same because the waiting time distribution calculation is not sensitive to the mode of degradation (first-order or delayed) since the absorbing state corresponds to the production of a new mature mRNA transcript or a new active Pol II which necessarily always precedes its degradation or removal. In addition to this reason, the statistics are the same for active Pol II and mature mRNA in the mechanistic model also because of the fixed time *τ* between the unpausing of a Pol II and the production of a mature mRNA.

Note that while the randomness parameter for *A* or *M* is greater than 1 for all parameter values (see (4)) in the two-state models, the same statistical measure can be less than or greater than 1 in the mechanistic model (see Eq. (5)). In fact, a graphical analysis of the latter equation shows that *R^mec^* ≥ 1/2; the 2 appears because in our model, it is the smallest number of reaction steps between *A* production events (*U*^⋆^ → *U*^⋆⋆^ → *U*^⋆^ + *A*). Similar results have been derived in the context of single molecule enzyme kinetics [48].

It follows that the two-state models can only capture the waiting time statistics of the mechanistic model (up to second order) when *R^mec^* ≥ 1 which is the case when the following condition is satisfied

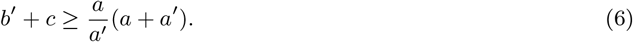

This implies that the conditions which favour a description of the mechanistic model by the two-state models are: (i) premature termination and unpausing from the paused promoter-proximal state must be fast i.e., large *b*′ + *c*; (ii) transcription factor binding to DNA elements and the reverse unbinding reaction must be slow i.e., small *a*+*a*′; (iii) transcription factor unbinding is fast compared to transcription factor binding i.e., *a/a*′ is small. Note that the condition gievn by Eq. (6) is not a function of *b*, the rate at which polymerase binds the promoter and moves to the proximal paused state (see later for an explanation of the role of *b*).

### 3.2 Analytical expressions for the effective parameters of the two-state models

Matching the first three moments of the waiting time distribution of the times between consecutive *M* or *A* production events of the telegraph/delayed telegraph model (given by Eqs. (27)) with those calculated using the mechanistic model (given by Eq. (40) evaluated for *i* = 1, 2, 3), we obtain a set of 3 simultaneous equations for the effective parameters of the two-state models *ρ, σ_u_, σ_b_*. The solution of these equations gives:

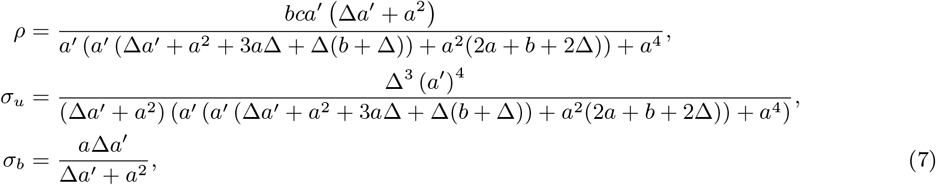

where Δ = *b*′ + *c − a* − (*a*^2^/*a*′). Note that if the condition given by Eq. (6) is satisfied, then Δ ≥ 0 and hence the effective parameters defined by Eqs. (7) are positive and physically meaningful. If the condition is not satisfied, then one of these effective parameters is negative which means that there are no two-state models that can approximate the mechanistic model’s waiting time moments up to third-order. We emphasize that the effective parameters are the same for the telegraph and delay telegraph models because the waiting time calculation is insensitive to the mode of decay (first-order or delayed). In Fig. 2(A-F) we compare the steady-state distribution of the two-state models evaluated with these effective parameters (for Δ > 0) and the steady-state distribution of the mechanistic model. In the cases shown, the two-state models provide an excellent match to the mechanistic model for both unimodal and bimodal distributions of active Pol II and mature mRNA numbers.

#### 3.2.1 The case of fast switching between *U*^⋆^ and *U*^⋆^

Where min(*b, b*′) ≫ max(*a, a*′), the two states *U*^⋆^ and *U*^⋆⋆^ can be effectively subsumed into a single super state *W* and the system dynamics amounts to switching between an inactive state *U* and an active state *W*. Physically, one sees that this arises since in this limit transitions between *U*^⋆^ and *U*^⋆⋆^ occur almost instantaneously compared to transitions between *U* and *U*^⋆^. The transition rate from *U* to *W* is the same as from *U* to *U*^⋆^ and hence in the two-state model this implies

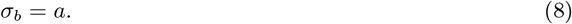

The transition rate from *W* to *U* must be equal to the transition rate from *U*^⋆^ to *U* multiplied by the probability of being in state *U*^⋆^ given that currently the effective system is in state *W*. This implies

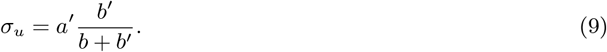

Similarly, the effective production rate is the rate of producing active Pol II from state *U*^⋆⋆^ multiplied by the probability of being in this state given that currently the effective system is in state *W*. This implies

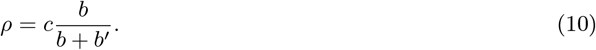

These results can be formally obtained from Eqs. (7) by choosing *b*′ = *γ_b_* (where *γ* is a constant) and taking the limit *b* → ∞. The case of fast switching in a similar three-state model of gene expression (without an explicit description of active Pol II dynamics) has been previously studied in [25]. While it is obvious that fast switching between *U*^⋆^ and *U*^⋆⋆^ simplifies to an effective two-state model, our condition (6) shows that fast switching is sufficient but not a necessary condition for a two-state model to describe the dynamics of the mechanistic model. We note that fast switching between *U*^⋆^ and *U*^⋆⋆^ is unlikely since the average time scale of Pol II pausing is ~7min [23] and almost 1 hour in a small subset of genes [50]. This indicates Pol II pausing is very stable and “not the consequence of fast, repeated rounds of initiation and termination” [23].

#### 3.2.2 Distinguishing between two-state and mechanistic models using waiting time distributions

It is interesting to note that while for Δ ≥ 0 the two-state and mechanistic models are practically indistinguishable by comparison of their number distributions, *they can be always distinguished by the distribution of the time between consecutive active Pol II or mRNA production events*. In particular, in Fig. 2(G-I) we show that while *f* (*t*), the waiting time distribution between consecutive production events, is a monotonically decreasing function for the two-state models, it has a peak at a non-zero value of time for the mechanistic model. Another distinguishing feature is that for two-state models, *f* (0) is non-zero while for the mechanistic model it is exactly zero. The latter feature can be explained as follows. For two-state models, since there is no change in the gene state when production occurs, hence there is no lower bound on how short the time between two consecutive production events can be. However, in the mechanistic model, a production event is accompanied by a change of state (from *U*^⋆⋆^ to *U*^⋆^), therefore there is a finite non-zero time to switch back to state *U*^⋆⋆^ from which the next production event occurs. Consequently, for the mechanistic model *f* (0) must be zero.

By this reasoning, it follows that the mode should be close to zero whenever the state *U*^⋆⋆^ is recovered rapidly after an active Pol II production event, which occurs when *b* is large. In Fig. 3 we confirm this intuition and show that for the mechanistic model as we increase *b*, the waiting time distribution of the two-state model better approximates the waiting time distribution of the mechanistic model. Note that a log-scale is used on the x-axis to emphasise that there are always differences between the mechanistic and two-state models for small values of *t*.

**Figure 3:**
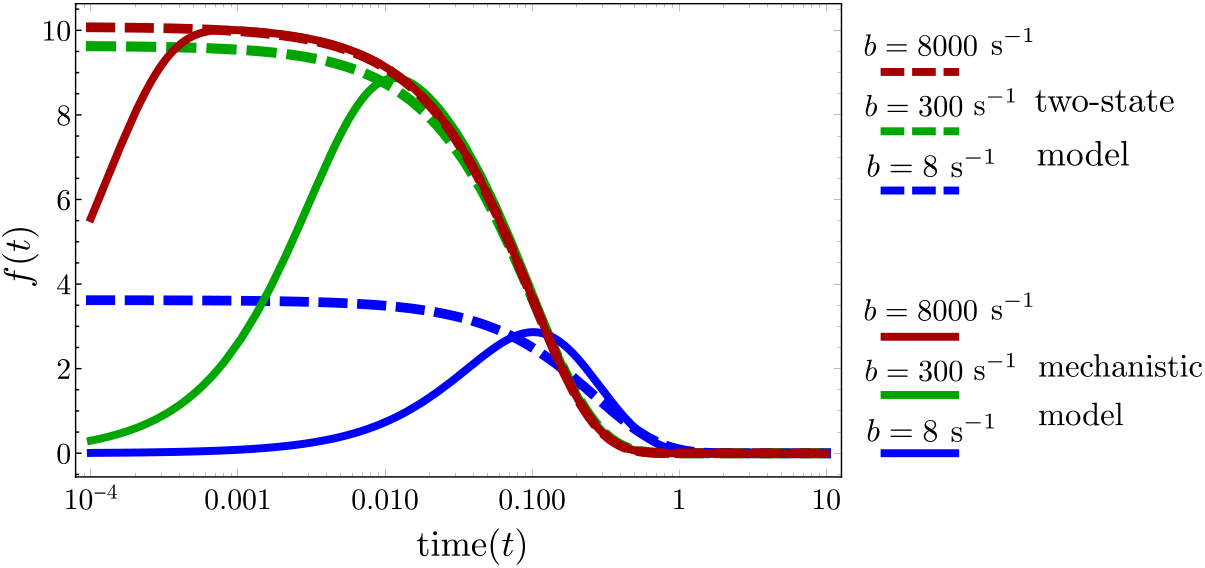
The waiting time distribution of the mechanistic model as a function of the rate parameter *b* (which controls the binding of Pol II to the promoter and the entry into the promoter-proximal state). The waiting time distribution of the mechanistic model is calculated by taking the inverse Laplace transform of Eq. (39). Note that as *b* increases, the peak moves closer to zero and the waiting time distribution of the mechanistic model approaches the waiting time distribution of the two-state model (calculated by taking the inverse Laplace transform of Eq. (26)). The parameter *b* is changed as described in the legend and the rest of the parameters are *a* = 0.1 s^−1^, *a*′ = 0.1 s^−1^, *b*′ = 4 s^−1^, *c* = 10 s^−1^. The parameters of the two-state model are calculated from Eqs. (7).

## 4 Sensitivity analysis

Equations (7) allow us to understand how the parameters of the mechanistic model influence the effective parameters of the two-state models. We define the ordered set of mechanistic model parameters as *θ^mec^* = {*a, a*′, *b, b*′, *c*} and the ordered set of the two-state model parameters as *θ^tele^* = {*ρ, σ_u_, σ_b_*}. In Table 1, we show the sign of the derivative of a parameter in a two-state model with respect to changes in the parameter of the mechanistic model (when Δ ≥ 0). For example, the first row shows the sign of the derivative of *ρ* with respect to *a, a*′, *b, b*′ and *c*. A positive (negative) sign for the pair (*ρ, a*) indicates that an increase in *a* leads to an increase (decrease) in *ρ*. We also show the same but for the burst size *β* = *ρ/σ_u_*, a commonly cited measure equal to the amount of mRNA produced while the gene is in the on state (in the two-state models). Note that while the sign is fixed for most cases, in three instances the sign can flip. There is also a case where one of the two-state model parameters is independent of one of the parameters of the mechanistic model (marked with a 0). Due to the complicated nature of Eqs. (7) it is difficult to deduce the signs in Table 1 using simple arguments, however in some cases it can be done. For example, the relationship of *ρ* with respect parameters *b, b*′ and *c* is intuitive since: (i) increasing *b* increases the time in the *U*^⋆⋆^ state meaning production of *A* (or *M* ) happens more often; (ii) decreasing *b*′ has the opposite effect, meaning production of *A* (or *M* ) occurs less often; and (iii) increasing *c* obviously increases the production rate of *A* (or *M* ) and hence increases the predicted value of *ρ*.

**Table 1:** Signs of the derivatives of the two-state effective parameters *ρ, σ_u_* and *σ_b_* with respect to the mechanistic model parameters *a, a*′*, b, b*′ and *c*. Expressions for the effective parameters are given by Eq. (7).

| | $a$ | $a'$ | $b$ | $b'$ | $c$ |
| --- | --- | --- | --- | --- | --- |
| $\rho$ | +/- | - | + | - | + |
| $\sigma_u$ | - | + | - | + | + |
| $\sigma_b$ | +/- | + | 0 | + | + |
| $\beta = \rho/\sigma_u$ | + | - | + | - | +/- |

Next, we investigate the sensitivities of the parameters *θ^tele^* of the two-state model to the parameters of the mechanistic model *θ^mec^*. For this purpose, we randomly selected 10^3^ parameter sets from a log-scaled space in the *θ^mec^* parameters, accepting only those parameter set combinations that came within 2 experimental errors of the measurements of Oct4 gene: *ρ* = 1.9±0.6 min^−1^, *σ_u_* = 1.8 × 10^−2^ ± 1.2 × 10^−2^ min^−1^ and *σ_b_* = 9.2 × 10^−3^ ± 2.8 × 10^−3^ min^−1^ [51]. We also did this for the Nanog gene whose measurements were: *ρ* = 0.8 ± 0.2 min^−1^, *σ_u_* = 6.9 × 10^−3^ ± 1.0 × 10^−3^ min^−1^ and *σ_b_* = 1.9 × 10^−3^ ± 0.2 × 10^−3^ min^−1^. A log-scaled parameter space was used such that various combinations of mechanistic model parameter timescales could be easily explored. The ranges of the mechanistic model parameters that we explored were 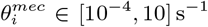 for the Oct4 gene and 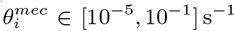 for the Nanog gene. The sensitivities calculated are the absolute values of the relative sensitivities given by,

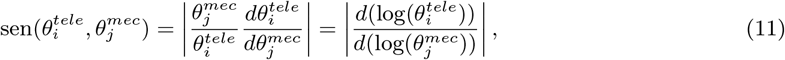

where 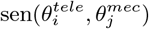 is the magnitude of the relative sensitivity of 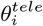 with respect to 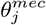.

As we show in Fig. 4, we find that for both genes, (i) the initiation rate *ρ* of the two-state models is most sensitive to parameters *b* and *c* in the mechanistic model i.e., parameters that control the rate of Pol II binding, of entering and leaving the promoter-proximal paused state; (ii) the rate of switching *off* of the two-state models *σ_u_* is most sensitive to parameter *a*′ (controlling transcription factor unbinding) and also to parameters *b, c* which control the initiation rate; (iii) the rate of switching *on* of the two-state models *σ_b_* is most sensitive to parameter *a* (controlling transcription factor binding) but also to parameters *a*′, *c* which control the the rate of switching *off* and the initiation rate. In Fig. 4, we also show which parameters of the mechanistic model are the three parameters of the two-state model least sensitive to. This analysis identifies how “microscopic” parameters of the mechanistic model affect the “macroscopic” parameters of the two-state models. *More importantly, it shows that the latter are typically correlated due to their dependence on common microscopic parameters.*

**Figure 4:**
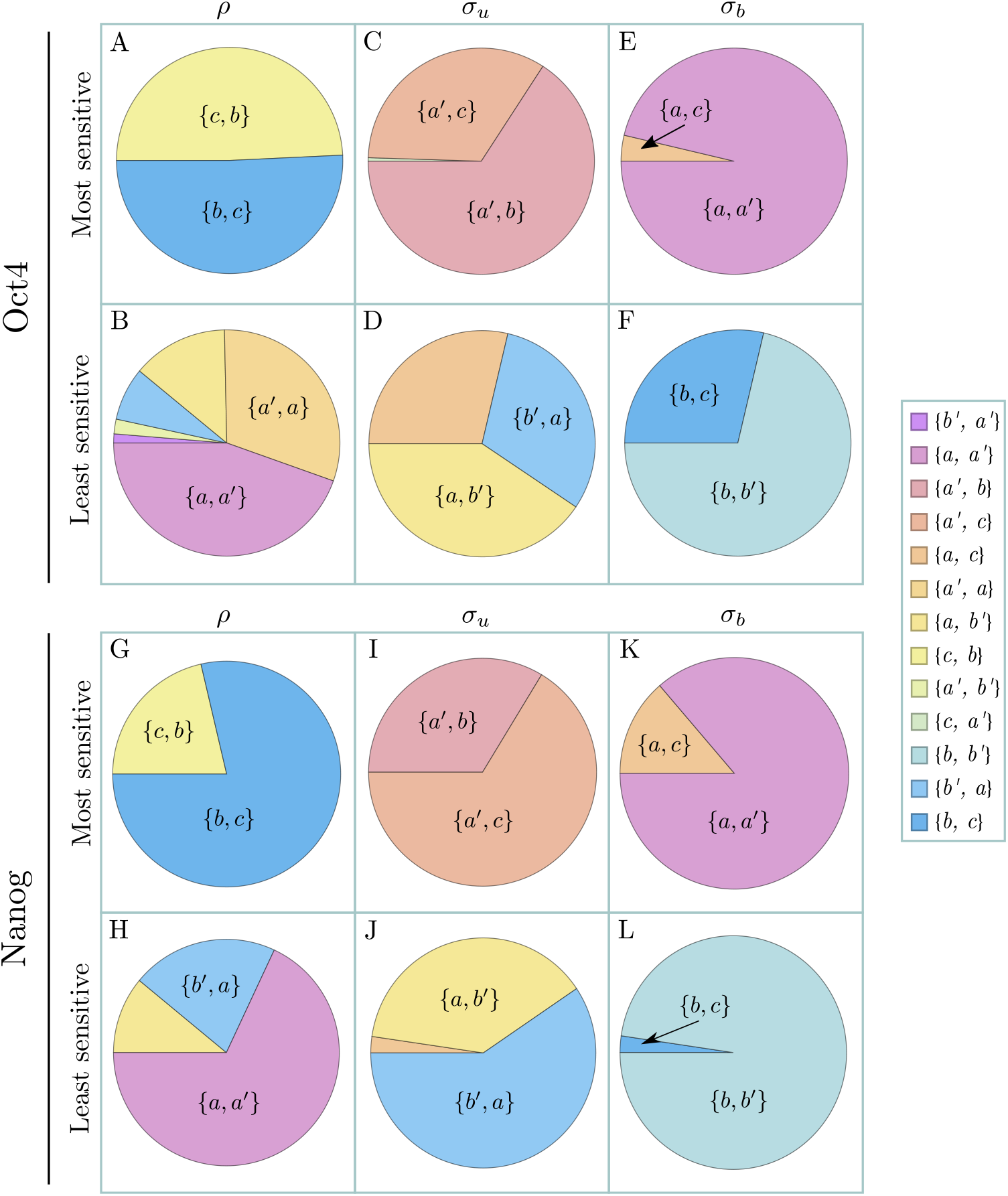
Pie charts showing the most and least sensitive of the telegraph model parameters *θ^tele^* = {*ρ, σ_u_, σ_b_*} with respect to mechanistic model parameters *θ^mec^* = {*a, a*′, *b, b*′, *c*}, for the Oct4 gene in (A-F) and for the Nanog gene in (G-L) [51]. We chose 10^3^ parameter sets *θ^mec^* at random, accepting only parameter sets for which the predicted telegraph model parameters *ρ, σ_u_* and *σ_b_* from Eq. (7) were within 2 experimental errors of values reported in [51]. From these randomly chosen parameter sets, we then calculated the relative sensitivity 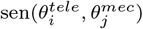 which is given by Eq. (11). The proportions on the pie charts show the proportion of parameter sets for which {*i, j*} were the most/least sensitive parameters, where {*i, j*} states that *i* is the most/least sensitive parameter followed by *j*. (A) for Oct4, the most sensitive parameters for *ρ* are *b* and *c*, with the majority of parameter sets being most sensitive to *b* and second-most to *c*. (B) for Oct4, the least sensitive parameters for *ρ* are *a* and *a*′, with the majority of parameter sets being least sensitive to *a* and second-least sensitive to *a*′. (C-F) follow similar interpretations as made for (A) and (B) for the Oct4 gene, and (G-L) follow similar interpretations for the Nanog gene.

## 5 Model reduction using number statistics or three-state models

Thus far, we have explored model reduction solely using waiting time statistics. Alternatively, one can match two-state and mechanistic models using moments of the number of molecules. As well, one can match three-state models and mechanistic models using waiting time or number statistics. In this section, we explore these alternative perspectives.

### 5.1 Obtaining reduced models with two states using number statistics

We begin by finding the Fano factors of the active Pol II and mature mRNA numbers in both the two-state and mechanistic models. In Appendices C and D, we derive expressions for the mean and variance of active Pol II and mature mRNA numbers in steady-state conditions for both the mechanistic and two-state models (for a test of their accuracy versus stochastic simulations using the delay SSA see Table 2). The Fano factor of the two-state models is easily proved to be always greater than 1. Specifically, for the delayed and standard telegraph models, we have respectively:

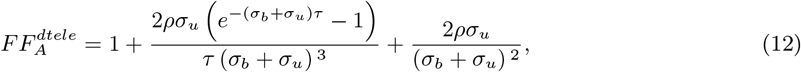

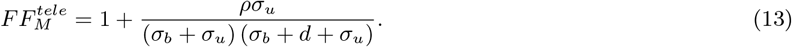

**Table 2:** Comparison of the mean and variance of *n* active Pol II and *m* mature mRNA numbers in the mechanistic model evaluated from the exact theory (Appendices C and D) and delay SSA (dSSA) with 10^5^ samples. 6 different parameter sets are considered.

| Method | $\langle n \rangle$ | $\text{Var}(n)$ | $\langle m \rangle$ | $\text{Var}(m)$ |
| --- | --- | --- | --- | --- |
| 1. $a = 0.016 \text{ s}^{-1}$, $a' = 0.08 \text{ s}^{-1}$, $b = 0.16 \text{ s}^{-1}$, $b' = 0.016 \text{ s}^{-1}$, $c = 0.24 \text{ s}^{-1}$, $d = 0.002 \text{ s}^{-1}$, $\tau = 273.62 \text{ s}$ | | | | |
| Theory | 6.195 | 17.554 | 14.151 | 27.701 |
| dSSA | 6.12 | 17.333 | 14.169 | 27.595 |
| 2. $a = 0.112 \text{ s}^{-1}$, $a' = 0.032 \text{ s}^{-1}$, $b = 0.16 \text{ s}^{-1}$, $b' = 0.016 \text{ s}^{-1}$, $c = 0.24 \text{ s}^{-1}$, $d = 0.016 \text{ s}^{-1}$, $\tau = 100 \text{ s}$ | | | | |
| Theory | 7.851 | 6.194 | 4.907 | 4.384 |
| dSSA | 7.664 | 6.08 | 4.908 | 4.38 |
| 3. $a = 0.144 \text{ s}^{-1}$, $a' = 0.032 \text{ s}^{-1}$, $b = 0.96 \text{ s}^{-1}$, $b' = 0.16 \text{ s}^{-1}$, $c = 0.24 \text{ s}^{-1}$, $d = 0.002 \text{ s}^{-1}$, $\tau = 273.62 \text{ s}$ | | | | |
| Theory | 43.511 | 37.646 | 99.387 | 92.745 |
| dSSA | 43.292 | 37.524 | 99.376 | 92.712 |
| 4. $a = 0.144 \text{ s}^{-1}$, $a' = 0.032 \text{ s}^{-1}$, $b = 1.12 \text{ s}^{-1}$, $b' = 0.8 \text{ s}^{-1}$, $c = 0.24 \text{ s}^{-1}$, $d = 0.1 \text{ s}^{-1}$, $\tau = 80 \text{ s}$ | | | | |
| Theory | 8.993 | 9.216 | 1.124 | 1.115 |
| dSSA | 8.904 | 9.127 | 1.124 | 1.114 |
| 5. $a = 0.032 \text{ s}^{-1}$, $a' = 0.032 \text{ s}^{-1}$, $b = 0.16 \text{ s}^{-1}$, $b' = 0.016 \text{ s}^{-1}$, $c = 0.32 \text{ s}^{-1}$, $d = 0.002 \text{ s}^{-1}$, $\tau = 273.62 \text{ s}$ | | | | |
| Theory | 16.838 | 36.098 | 38.462 | 61.715 |
| dSSA | 16.707 | 35.825 | 38.473 | 61.668 |
| 6. $a = 0.016 \text{ s}^{-1}$, $a' = 0.032 \text{ s}^{-1}$, $b = 0.16 \text{ s}^{-1}$, $b' = 0.016 \text{ s}^{-1}$, $c = 0.4 \text{ s}^{-1}$, $d = 0.005 \text{ s}^{-1}$, $\tau = 50 \text{ s}$ | | | | |
| Theory | 2.273 | 5.909 | 9.091 | 21.629 |
| dSSA | 2.185 | 5.6 | 9.096 | 21.611 |

The Fano factor of the number of active Pol II in the mechanistic model is given by:

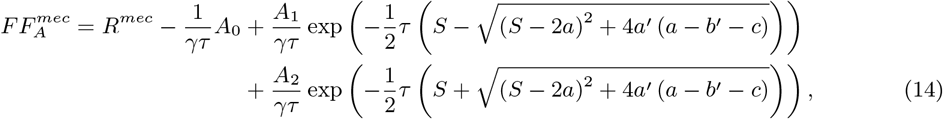

where *A*_0_, *A*_1_ and *A*_2_ can be positive or negative and are complicated functions of *a, a*′, *b, b*′, *c*, and where we have defined

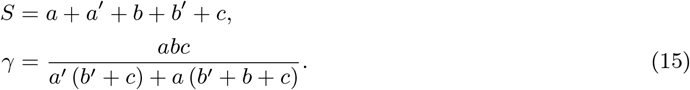

Note that the arguments of the exponential functions are negative for all positive values of the parameters. The Fano factor of the mature mRNA statistics is derived in Appendix D and is given by:

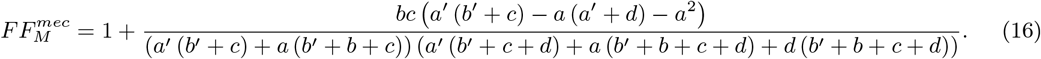

*Since* 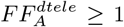 *and* 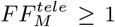, *clearly model reduction using molecule number moments will only be possible if the parameters of the mechanistic model are such that* 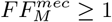 *and* 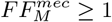. For the mature mRNA, this analysis is straightforward. Similar to the derivation of the condition (6), from the numerator of the second term in Eq. (16), it can be deduced that 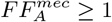 provided the following condition holds:

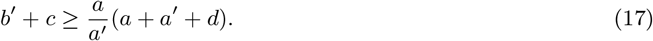

When condition (17) is satisfied, one can find a mapping between the standard telegraph model describing mature mRNA and the mechanistic model. We note that this condition is not the same as that derived from model reduction using waiting time statistics, namely Eq. (6). In fact, if Eq. (6) is not satisfied then neither is Eq. (17) i.e., for all points in parameter space in which it is not possible to match the moments of waiting time distributions of the two-state and mechanistic models, it is also not possible to match the moments of the mature mRNA numbers. However, it also follows that there is a region of parameter space of the mechanistic model where moment matching of the two-state model using waiting time statistics is possible (Eq. (6) is satisfied) whereas moment matching using number statistics is not (Eq. (17) is not satisfied). This region of parameter space where the two methods give different results is very small whenever *a*+*a*′ ≫ *d* where the rates of transcriptional factor binding/unbinding to the promoter are much larger than the rate of mature mRNA degradation. This seems to be generally the case since degradation timescales of mature mRNA are generally many hours in eukaryotic cells [8]. Incidentally, this offers an explanation why the Fano factor of mature mRNA is invariably measured to be greater than 1 in the literature of eukaryotic gene expression [9, 10, 52].

Due to the complicated nature of the expression in Eq. (14), the derivation of an analytic condition for which the Fano factor of active Pol II is greater than 1 appears to be difficult to obtain. However, in the limit of *τ* → ∞ it is clear that 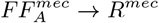. Hence, in the limit of long elongation times, the condition necessary for model reduction using active Pol II moment number statistics i.e., 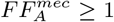, is equivalent to the condition necessary for model reduction using waiting time statistics given by Eq. (6) (which is the same as *R^mec^* ≥ 1). This is intuitive since the waiting time calculation does not consider the removal of active Pol II via elongation but only their production time statistics. It can also be proved from Eq. (14) that in the limit *τ* → 0 we have 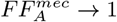. What happens for finite *τ* > 0 is difficult to deduce from Eq. (14) and hence we investigate this numerically.

In Fig. 5 we evaluate Eq. (14) as a function of *τ* for a number of parameter sets with different *R^mec^*. Several notable features can be seen: (i) if *R^mec^* < 1 then 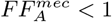 i.e., if model reduction using waiting time statistics is not possible then it is also impossible using number statistics; (ii) for *R^mec^* ≥ 1, as we increase *τ*, 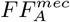 decreases from 1 to a value below 1, reaches a minimum and then increases up to the value *R^mec^*. *Consequently, if the condition for model reduction using waiting time statistics is satisfied, it is not necessarily true that it is possible to achieve model reduction according to number statistics.* In Fig. 6 (A-C), we show a heat map of the minimum Fano factor (achieved at intermediate *τ*) in the parameter space of the mechanistic model. Note that the minimum achieved inside the region where *R^mec^* > 1 (the region above the white line) is not far below 1. As a consequence, while here there is no model reduction from a number statistics point of view, model reduction using waiting time statistics is possible, and the distribution computed using the effective parameters given by Eqs. (7) while not perfect, it is acceptable – see Fig. 6 (D-F).

**Figure 5:**
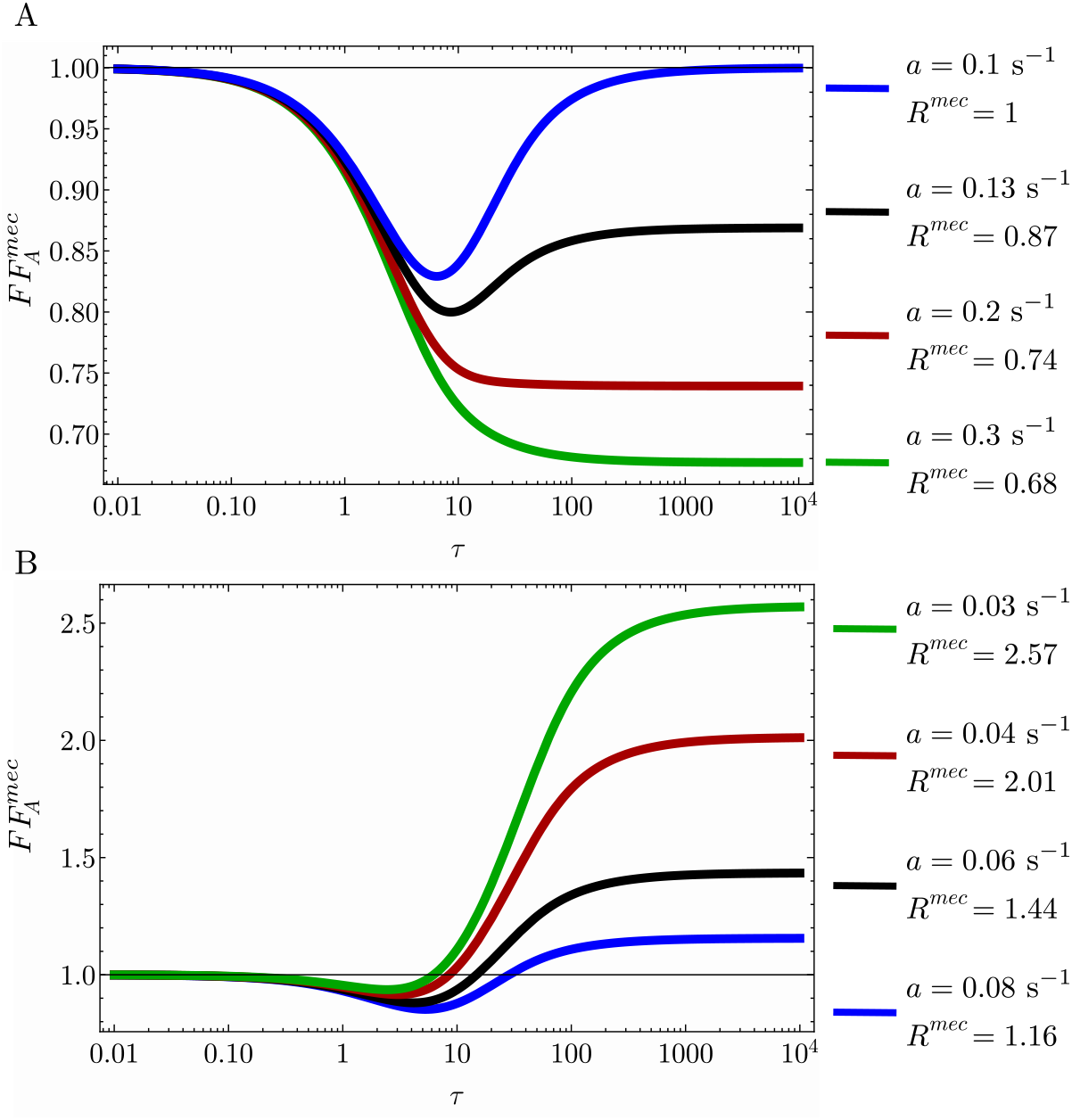
Fano factor of the active Pol II number distribution for the mechanistic model as a function of the elongation time *τ* and the randomness parameter *R^mec^*. The Fano factor is evaluated using Eq. (14). Note that the large *τ* limit of the Fano factor is equal to the randomness parameter *R^mec^* which is given by Eq. (5); *R^mec^* is here varied via the parameter *a* whilst keeping the rest of parameters constant: *b*′ = 0.0125 s^−1^, *a*′ = 0.032 s^−1^, *b* = 0.16 s^−1^, *c* = 0.4 s^−1^. (A) shows that if *R^mec^* ≤ 1 then the Fano factor is less than 1 for all *τ*. (B) shows that if *R^mec^* ≥ 1 then the Fano factor is less than 1 for a small enough value of *τ*.

**Figure 6:**
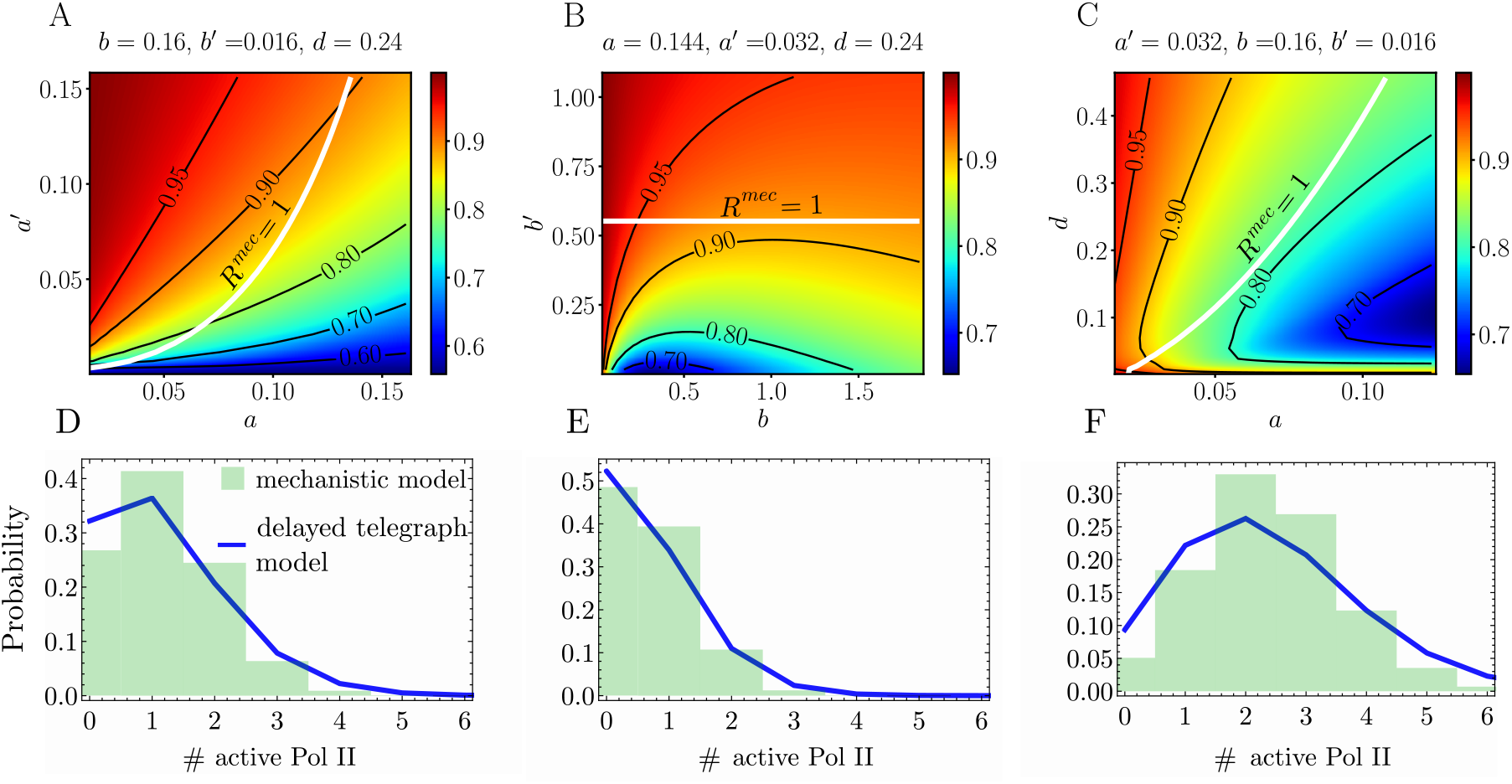
Accuracy of the number distributions of active Pol II constructed using the delay telegraph model for parameters close to the boundary *R^mec^* = 1. (A-C) The minimum Fano factor of active Pol II numbers in the mechanistic model as a function of the parameters *θ^mec^* = {*a, a*′, *b, b*′, *c*}. For a given set of *θ^mec^*, the minimum Fano factor is found by varying the elongation time *τ* in Eq. (14). The region above the white line *R^mec^* = 1 is where model reduction using waiting time statistics is possible. Note that inside this region, the minimum Fano factor of active Pol II numbers is smaller than 1 meaning that for small enough values of *τ*, model reduction using number statistics is not possible. However, in (D-F) we show that even when this is the case, the number distributions constructed using the delay telegraph model with effective parameters given by Eq. (7) provide a reasonably good approximation to the mechanistic model distribution of active Pol II. Note that these distributions represent a worst case scenario – inside the boundary, for the vast majority of points, the two-state model distributions provide an almost perfect fit to the mechanistic model distributions as shown in Fig. 2. The parameters are as follows: (D) *a* = 0.0336 s^−1^, *a*′ = 0.006 s^−1^, *b* = 0.16 s^−1^, *b*′ = 0.016 s^−1^, *c* = 0.24 s^−1^, *τ* = 13.65 s, 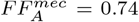, *R^mec^* = 1.034. (E) *a* = 0.072 s^−1^, *a*′ = 0.0288 s^−1^, *b* = 0.16 s^−1^, *b*′ = 0.016 s^−1^, *c* = 0.24 s^−1^, *τ* = 8.76 s, 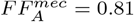, *R^mec^* = 1.004. (F) *a* = 0.08 s^−1^, *a*′ = 0.005 s^−1^, *b* = 2 s^−1^, *b*′ = 0.0001 s^−1^, *c* = 1.36 s^−1^, *τ* = 3 s, 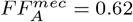, *R^mec^* = 1.0001.

Thus far, we have looked at model reduction via number statistics from the perspective of when the Fano factor numbers of the mechanistic and two-state models are both greater than one. In Appendix E we extend this analysis further by considering two other types of model reduction via number statistics: (1) matching of the molecule number moments and (2) of the number distributions of the mechanistic and the two-state models for active Pol II and mature mRNA numbers. In particular, we found the following: (i) within the region of parameter space of the mechanistic model described by the condition Eq. (6), it was possible to numerically find parameters of the two-state models such that the first three moments of the active Pol II and mature mRNA number distributions of the two-state models agreed with those of the mechanistic model – see Fig. 7 (A-C) and (G-I); (ii) the Hellinger distance between the molecule number distributions predicted by the mechanistic model and the molecule number distributions of the two-state models that provides the best approximate distribution of the mechanistic model, is very small within the region defined by Eq. (6) – see Fig. 7 (D-F) and (J-L). The analysis shows there is a close relationship between model reduction using waiting time and number statistics, and supports the conclusions reached in Sections 3 and 4 using waiting time statistics.

**Figure 7:**
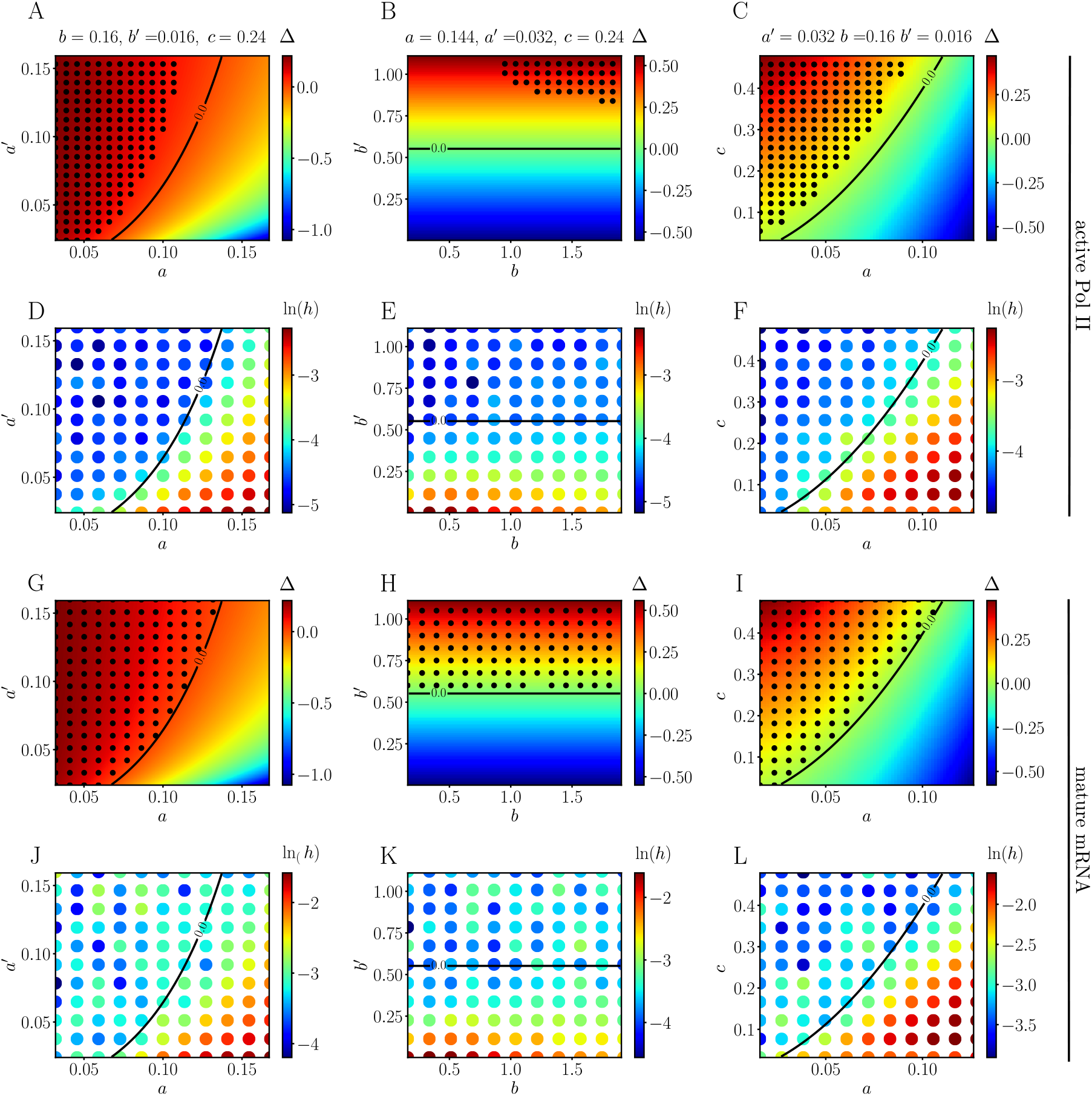
Comparison of model reduction of the mechanistic model to two-state models using two different types of number statistics and comparison with reduction from waiting time statistics. In (A-C) black dots show the points in parameter space where the first 3 moments of the active Pol II number distributions of the mechanistic and the delayed telegraph model match numerically using the Newton-Raphson method; in (G-I) we show the same for the distributions of mature mRNA of the mechanistic and telegraph models. The heat map shows the the value of Δ = *b*′ +*c − a* − (*a*^2^/*a*′) and the solid black lines divides areas where Δ > 0 (waiting time moment matching exists) and Δ < 0 (waiting time moment matching does not exist). Note that the black dots in A-C do not fill the whole region Δ > 0 because of numerical issues with the solver (See Appendix E for a discussion). In (D-F) and (J-L) we show the Hellinger distance (*h* in log scale) between the molecule number distributions predicted by the mechanistic model and the molecule number distributions of the two-state models that provides the best approximate distribution of the mechanistic model; the parameters of the two-state models are those learnt after *O*(10^5^) iterations of an algorithm that maximizes the likelihood. The mature mRNA decay rate *d* = 0.0016 s ^−1^ in all cases and the delay time is *τ* = 273.62 s. See Appendix E for details of the numerical procedures used.

### 5.2 Obtaining reduced models with three states using waiting time statistics

Thus far, we have considered the approximation of the mechanistic model by two-state models (telegraph and delay telegraph models). However, some papers have postulated the existence of two off states for some mammalian genes because the time spent by the gene in the off state is measured to be non-exponential [11]. This has led to a variation of the telegraph model, which we will refer to as the refractory model

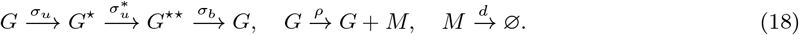

One can also postulate a modification (parallel to the delay telegraph model) that describes active Pol II rather than mature mRNA:

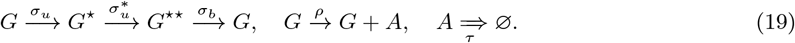

An analysis akin to the one shown for the two-state models in Appendix A.1 shows that the Laplace transform of the waiting time distribution of the time between consecutive active Pol II or mature mRNA production events is given by:

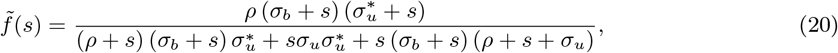

where 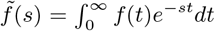. From the definition of the Laplace transform, we have that the moments are given by

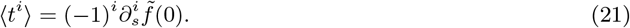

The randomness parameter is then given by the square of the coefficient of variation squared of the time between two consecutive production events:

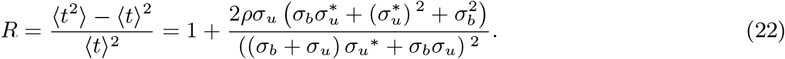

Hence the randomness parameter of the reduced models (18) and (19) is always greater than 1. In contrast, we have already shown by Eq. (5) that for the mechanistic model the randomness parameter can be greater than or less than 1. Hence it follows that the condition given by Eq. (6) is necessary for both telegraph models and those with a refractory state to approximate the mechanistic model. Similar to what we have previously done for the two-state models, analytical expressions expressing the four parameters of the reduced refractory models in terms of the six parameters of the mechanistic model can be derived by matching the first four moments of the waiting time distribution of the two models. The steady-state distribution solutions of active Pol II and mature mRNA numbers of the reduced refractory models evaluated with these effective parameters provide an excellent approximation to the distributions of the mechanistic model. However, since this was already achieved using two-state models and since the refractory models have the same limitations as the two-state models (randomness parameter cannot be less than 1), it follows that *two-state models provide the optimal choice for model reduction within the parameter space defined by* Eq. (6).

## 6 Discussion

In this paper we have investigated to what extent can two-state models predict the active Pol II and mature mRNA dynamics of a more realistic mechanistic model that incorporates transcriptional factor binding and unbinding, Pol II dynamics (binding, pausing, release, elongation) and mature mRNA dynamics. We found that there is a region of parameter space where there exists a choice of parameters of two-state models in terms of the mechanistic model such that the first three moments of their waiting time distributions exactly match. The distributions of active Pol II and mature mRNA numbers predicted by two-state models with these effective parameters provide a very close match to the distributions predicted by the mechanistic model; nevertheless, the models can be distinguished by comparison of the shape of their waiting time distribution. The waiting time distribution for the two-state model has a non-zero value at *t* = 0 and decreases monotonically with time; whereas for the mechanistic model, the waiting time distribution is zero at *t* = 0 and has a peak at a non-zero value of time. We note that while in principle these two distributions are always distinguishable, in practice the differences will be small if the rate of Pol II binding and entry into the paused state is very large. We also showed that the necessary condition for the reduction of the mechanistic model to two-state models that was analytically derived using waiting time statistics i.e., Eq. (6), is compatible with the region of parameter space identified by model reduction using matching of moments and distributions of molecule numbers. We note that while our model description was framed in terms of an activator, it has alternative interpretations which increase its generality and applicability. One such alternative interpretation is in terms of a repressor that operates via competitive binding [53, 54]. In this interpretation *U* is a state that has a repressor bound to the promoter such that Pol II is blocked from being able to bind to the promoter. *U*^⋆^ then represents the state where the promoter is free and neither repressor nor Pol II is bound to the promoter, meaning that the binding site is accessible to both repressors and Pol II. Finally, the *U*^⋆⋆^ state represents the state in which Pol II is recruited and proximally-paused.

A main distinction of this work from the analysis of a similar model studied in [25] is that the present mechanistic model has an explicit description of active Pol II that allows us to study the accuracy of the delay telegraph model. It is also noteworthy that while [25] showed that the telegraph model provided an excellent approximation to the mature mRNA distribution of a similar mechanistic model under the assumption of rapid entry and exit from the paused state, in this study we showed using a variety of model reduction techniques that this assumption though sufficient is not necessary. We also note that while other papers have made use of waiting time statistics in the context of gene expression [11], our approach is distinctly different. The distribution of the off time in another three-state model of gene expression [11] is not the same as the distribution of the time between two consecutive active Pol II production events; this is because the former provides only information about the time between two successive bursts of gene expression which occurs on long timescales and reflects the accessibility of the promoter but has no information on the fast Pol II processes within each burst. To the best of our knowledge, the experimental measurement of the distribution of the waiting time as defined in our paper has not been attempted yet. This is because with current labelling and imaging technology, it is not easy to directly visualize, track and quantify individual transcriptional initiation events. However a set of recent papers report progress in this direction by estimating an approximate distribution between two consecutive initiation events in Drosophila using a machine-learning approach [55, 56].

We finish by a discussion of the validity and interpretation of Eqs. (7) which express the parameters of two-state models as a function of the parameters of the mechanistic model. We have shown from these expressions that different parameters of the two-state models can be effectively correlated due to their dependence on a common parameter/s of the mechanistic model. This may explain correlations found between parameters of two-state models estimated from single cell RNA sequencing for mammalian cells [10]. There is a region of parameter space where the effective parameters given by our theory become negative (when the inequality given by Eq. (6) is not satisfied), meaning that in this case there is no two-state model that can match the first three moments of the waiting time distribution of the mechanistic model; we also showed that if the elongation time and the mature mRNA degradation timescale are large enough, the aforementioned region is also characterized by Fano factors of active Pol II numbers and mature mRNA numbers that are less than one i.e., sub-Poissonian statistics. To see whether such a case is realistic we extensively searched through the experimental literature of gene expression, and found that for mature mRNA all papers report a Fano factor of greater than 1 which is consistent with constitutive or bursty expression; for nascent RNA, the majority of papers reports Fano factors greater than 1 (see for example [9, 20, 57]) with the exception of one paper (see Supplementary Fig. 6 of [58]). However, it is to be borne in mind that while theoretically nascent RNA numbers should equal the active Pol II numbers in our model, in practice due to the intricacies of smFISH this is not the case, as we now explain. The number of nascent mRNA is most commonly calculated by dividing the total fluorescent signal from a transcription site by the fluorescence emitted by a mature transcript. In this technique, a fluorescent signal is emitted by oligonucleotide probes bound to the nascent RNA tail. Since as an active Pol II travels along the gene, its nascent RNA tail grows, we expect the fluorescent signal intensity to increase as well [14]. Hence it follows that the total nascent mRNA *n_N_* calculated using this method is generally a lower bound on the actual number of active Pol II *n_A_* at a transcription site in the nucleus i.e., *n_N_* ~ *fn_A_* where *f* is a fraction. From this it immediately follows that the Fano factor of nascent mRNA is always less than the Fano factor of active Pol II. Thus the measurement of Fano factors of nascent mRNA numbers slightly less than 1 in [58] likely implies Fano factors of active Pol II which are above 1. Hence we come to the conclusion that all available evidence to date for both nascent and mature mRNA seems consistent with Eq. (6), which implies that Eq. (7) provides a generally useful means to understand the parameters of two-state models in terms of underlying microscopic processes.

## Acknowledgments

S. B and R. G were supported by a Leverhulme Trust grant (RPG-2018-423). J.H was supported by a BBSRC EASTBIO PhD studentship.

## A Waiting time calculations for two-state models

## A.1 Derivation of the waiting time distribution and its moments

Consider the delayed telegraph model describing active Pol II dynamics:

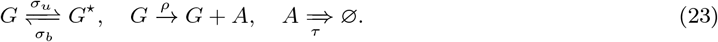

We want to calculate the distribution of the waiting time between the production of two consecutive active Pol II molecules (*A*) along the gene. In other words, given that a paused Pol II has just been released and become active, what is the distribution of the time before the next Pol II becomes active? Note that the mechanism of removal of active Pol II does not influence the statistics of the production events. Hence the calculation that proceeds remains the same if instead of the delayed telegraph model, we had to use the telegraph model to calculate the waiting time distribution between two consecutive mature mRNAs.

We define three states: state *X* where the gene is in state *G* and the number of active Pol II is *n*; state *Y* where the gene is in state *G*^⋆^ and the number of active Pol II is *n*; state *Z* where the gene is in state *G* and the number of active Pol II is *n* + 1. Hence the effective reaction scheme describing these three states is

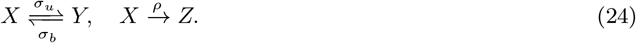

Immediately after an active molecule of Pol II is produced, the gene is in state *G* and hence our initial condition is state *X*. The absorbing state is state *Z*. The master equations describing the effective reaction scheme are:

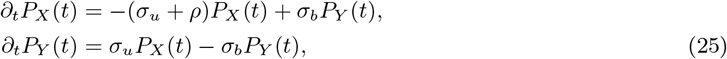

with initial condition *P_X_* (0) = 1, *P_Y_* (0) = 0 and *P_Z_* (0) = 0. The distribution *f* (*t*) of the time *t* at which the system enters the absorbing state *Z* is given by the probability that the system is in state *X* at time *t* multiplied by the rate of switching from state *X* to *Z*, i.e. *f* (*t*) = *ρP_X_* (*t*). Solving the differential equations Eq. (25) using the Laplace transform we obtain:

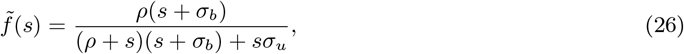

where 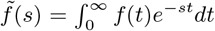. It then follows that the first three moments of the time between two consecutive active Pol II production events are given by:

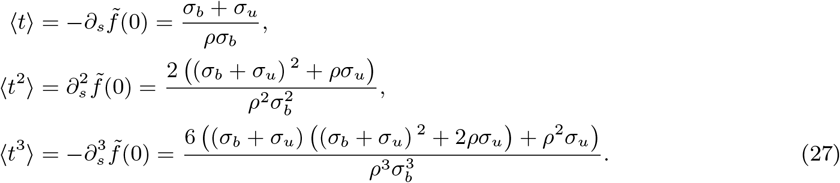

The square of the coefficient of variation of the waiting time (the randomness parameter) is:

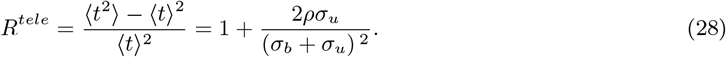

Note that *R^tele^* > 1 for all parameter values. For reference, the exponential distribution is characterized by a coefficient of variation squared equal to 1.

## A.2 Proof of the monotonicity of the waiting time distribution

Here we prove that the waiting time distribution of the delayed telegraph model (and of the telegraph model) is a monotonically decreasing function. We start by rewriting Eq. (26) in the form:

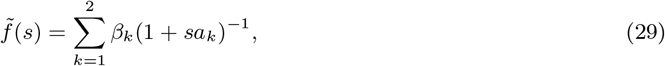

where

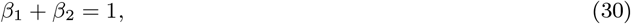

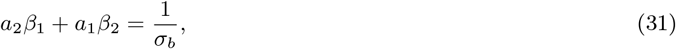

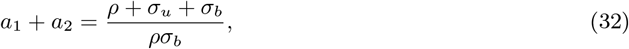

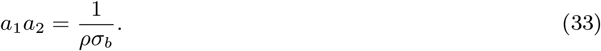

Taking the inverse Laplace transform of Eq. (29) one can show that

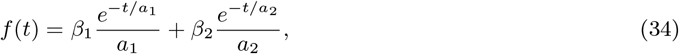

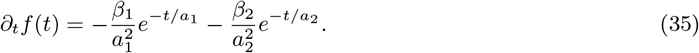

To determine if *f* (*t*) is monotonically decreasing in *t*, we need to know what is the sign of *a*_1_, *a*_2_, *β*_1_, *β*_2_. From Eqs. (32) and (33), since the right hand sides of both equations are positive then *a*_1_, *a*_2_ must also be positive (if one or both are negative then the sign of the left hand side will not match the sign on the right hand side of one of the two equations). Also by solving Eqs. (30) and (31) simultaneously for *β*_1,2_ one finds that these are positive. Because *a*_1_, *a*_2_, *β*_1_, *β*_2_ > 0, it follows from Eq. (35) that *∂_t_f* (*t*) < 0 for all times and hence *f* (*t*) is a monotonic decreasing function of time *t*. Furthermore, by the initial value theorem [59] and Eq. (26), we have 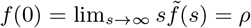.

## B Waiting time calculations for the mechanistic model

## B.1 Derivation of the waiting time distribution and its moments

In this section, we extend the analysis of Appendix A to study the mechanistic model, which is given by:

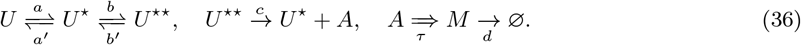

We now derive the distribution of the time between two consecutive active Pol II production events and also the same but for mature mRNA *M*. We first consider the active Pol II case. We define four states: state *W* where the gene is in state *U* and the number of active Pol II is *n*; state *X* where the gene is in state *U*^⋆^ and the number of active Pol II is *n*; state *Y* where the gene is in state *U*^⋆⋆^ and the number of active Pol II is *n*; state *Z* where the gene is in state *U*^⋆^ and the number of active Pol II is *n* + 1. Hence the effective reaction scheme describing these four states is

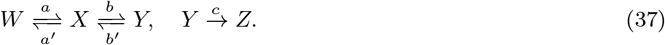

Just after an active Pol II is produced, the gene is in state *U*^⋆^ and hence our initial condition is state *X*. The absorbing state is state *Z*. The master equations describing the effective reaction scheme are:

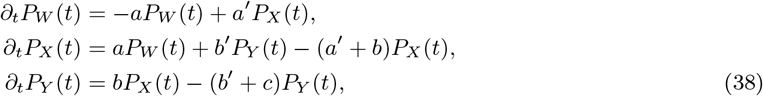

with initial condition *P_X_* (0) = 1, *P_W_* (0) = *P_Y_* (0) = *P_Z_* (0) = 0. The distribution *f* (*t*) of the time *t* at which the system enters the absorbing state *Z* is given by the probability that the system is in state *Y* at time *t* multiplied by the rate of switching from state *Y* to *Z*, i.e. *f* (*t*) = *cP_Y_* (*t*). Solving the differential equations Eq. (38) using the Laplace transform we obtain:

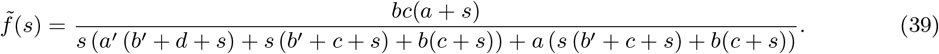

From the definition of the Laplace transform, we have that the moments are given by

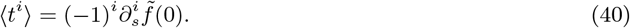

The square of the coefficient of variation squared of the time between two consecutive production events (the randomness parameter) is:

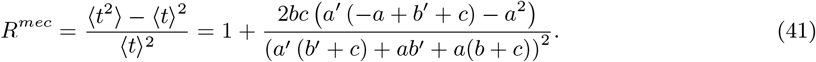

Note that depending on the parameter values, *R^mec^* can be greater than or less than one (unlike for two-state models where it was shown in Appendix A that the randomness parameter is always greater than one).

Suppose there is a fixed time *τ* between the production of an active Pol II and the production of a mature mRNA (via elongation and termination). It follows that the time between two consecutive mature mRNA production events is precisely the same as the time between two consecutive Pol II activation events, i.e. all the waiting time statistics that we have derived for active Pol II also hold for mature mRNA too.

## B.2 Some properties of the waiting time distribution

We note that since 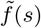 in Eq. (39) can be written in the form 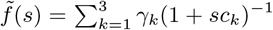 (for particular values of the constants *γ_k_* and *c_k_*), it follows that

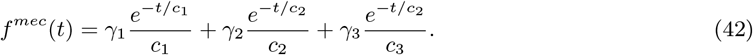

This is unlike that for two-state models in Appendix A where the waiting time distribution was a sum of two exponentials.

Also by the initial value theorem and Eq. (39), we have that 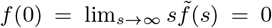. As well necessarily for any distribution we have that lim_*t*→∞_ *f* (*t*) = 0. Hence it follows by the behavior of *f* (*t*) at *t* = 0 and *t* = ∞, that the positive function *f* (*t*) must achieve one or more maxima at intermediate times. Hence the waiting time distribution for the mechanistic model is non-monotonic in time *t* (unlike for two-state models, which have a monotonic waiting time distribution).

## C Derivation of the steady-state mean and variance of active Pol II numbers for the mechanistic model

We first calculate the statistics of the accumulated active Pol II on the gene, i.e. ignoring its removal due to elongation. Hence we want to derive the time-dependent first and second moments of the reaction scheme:

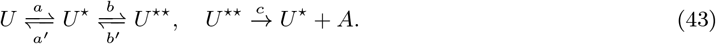

The easiest way to calculate these moments is using *the linear-noise approximation, which is exact up to second-order moments for any system with linear propensities (as in our case)*. The stoichiometric matrix and the propensity (column) vector are given by:

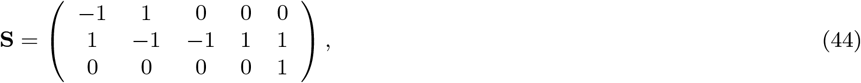

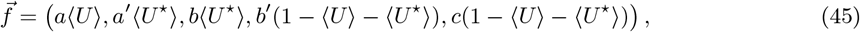

where 〈*ψ*〉 denotes the average number of molecules of species *ψ*. The species are numbered in the order *U*, *U*^⋆^, *A* and the reactions in the order *U* → *U*^⋆^, *U*^⋆^ → *U*, *U*^⋆^ → *U*^⋆⋆^, *U*^⋆⋆^ → *U*^⋆^, *U*^⋆⋆^ → *U*^⋆^ + *A*. The matrix element [**S**]_*ij*_ is the net change in the number of molecules of species *i* when reaction *j* occurs, and the vector element *f_j_* is the average propensity of the *j^th^* reaction. Note that we have used the conservation law 〈*U*^⋆⋆^〉 = 1 − 〈*U*〉 − 〈*U*^⋆^〉 to simplify the vector 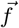.

The equations for the first two moments are given by:

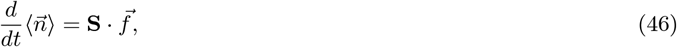

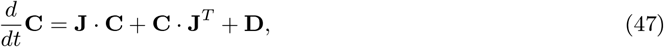

where 〈*n_i_*〉 is the average number of molecules of species *i* and [**C**]_*ij*_ = *C_ij_* is the covariance between species *i* and *j*. Furthermore we have defined the matrix **J** as the Jacobian of the rate equations Eq. (46) and **D** as the diffusion matrix which equals 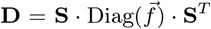. The matrix 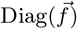 is a diagonal matrix with diagonal elements given by the elements of the vector 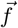.

The time-dependent solution of these equations is quite complex since we have three interacting species. However, the calculation is much simplified if one makes use of the fact that *U*, *U*^⋆^, *U*^⋆⋆^ will reach a steady-state after some time. This implies that

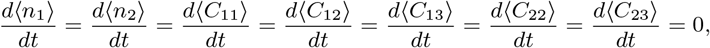

which leads to the solutions:

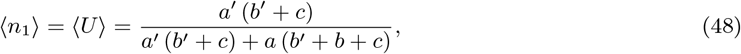

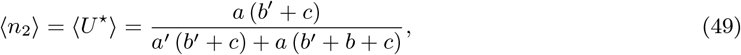

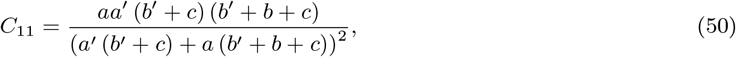

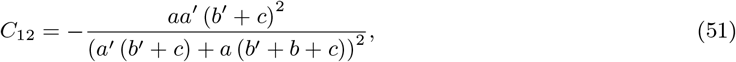

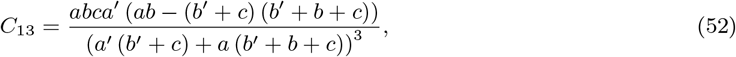

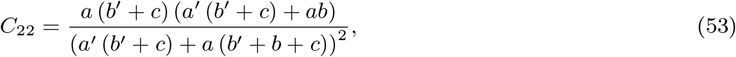

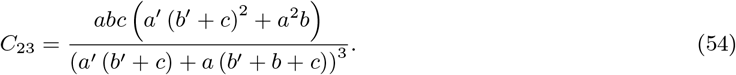

However, since active Pol II keeps accumulating with time, we have to solve the time-dependent equations for its mean and variance which from Eqs. (46) and (57) are given by:

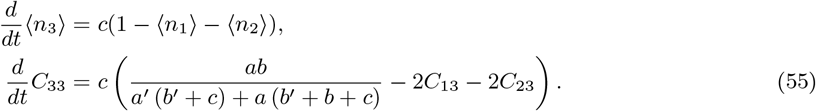

Substituting Eqs. (48), (49), (52) and (54) in Eq. (55) and solving the resulting differential equations with zero initial conditions, we finally obtain the time-dependent mean and variance of the accumulated active Pol II:

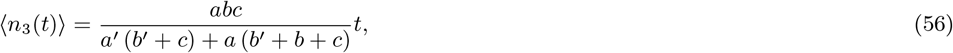

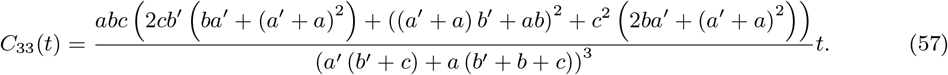

Hence the Fano factor of accumulated active Pol II is given by:

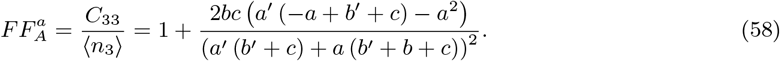

Note that 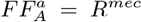 given by Eq. (5). This equivalence between the Fano factor of accumulated products and the coefficient of variation of the waiting times has been previously reported in the single enzyme molecule literature [48].

Next we use these results to calculate the mean and variance of active Pol II in steady-state conditions, i.e., the statistics of active Pol II due to both binding and unbinding reactions. Let the number of observed active Pol II at time *t* be *n*(*t*); then it follows that if elongation happens after a deterministic time *τ* we can write:

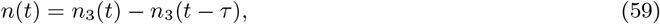

where *n*_3_(*t*) is the number of active Pol II accumulated up till time *t*. This relationship between the observed number of active Pol II and the number of accumulated active Pol II follows from the elongation dynamics: since all active Pol II molecules have a fixed lifetime of *τ*, it follows that molecules produced before time *t − τ* must have all died by time *t* and only those produced in the interval (*t − τ, t*] will contribute to the number of observed molecules at time *t*. Hence the first two moments of the observed active Pol II at time *t* are given by:

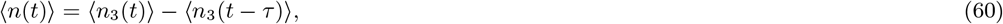

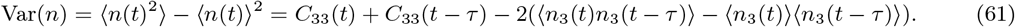

The equation for the steady-state mean Eq. (60) can be easily evaluated by means of Eq. (56) leading to:

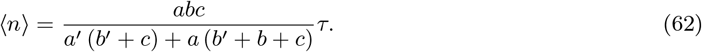

To calculate the steady-state variance of observed active Pol II, we need to first evaluate the correlator 〈*n*_3_(*t*)*n*_3_(*t − τ*)〉 − 〈*n*_3_(*t*)〉 〈*n*_3_(*t − τ*)〉 which appears on the right-hand side of Eq. (61). Following Gardiner [60], for any linear system, the autocorrelation vector in steady-state conditions 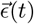 with elements

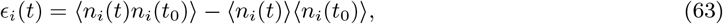

obeys the differential equation:

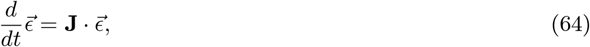

with the initial condition given by **C**(*t* = *t*_0_). Hence we have

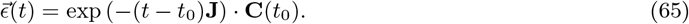

Choose *t*_0_ = *t* − *τ*, it follows that the correlator 〈*n*_3_(*t*)*n*_3_(*t* − *τ*)〉 − 〈*n*_3_(*t*)〉〈*n*_3_(*t* − *τ*)〉 is equal to *ε*_3_(*t*). Note that **C**(*t*_0_) = **C**(*t* − *τ*) has elements given Eqs. (50)–(54) and (57). Hence we can finally evaluate Eq. (61):

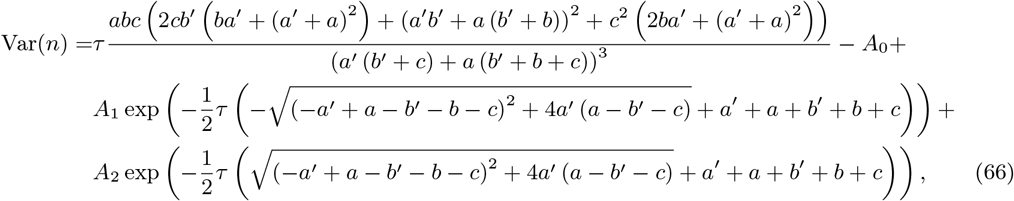

where

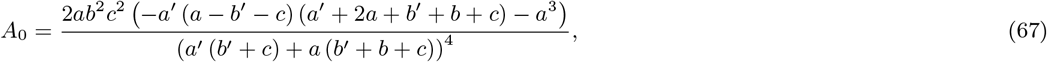

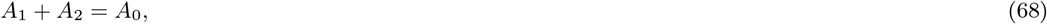

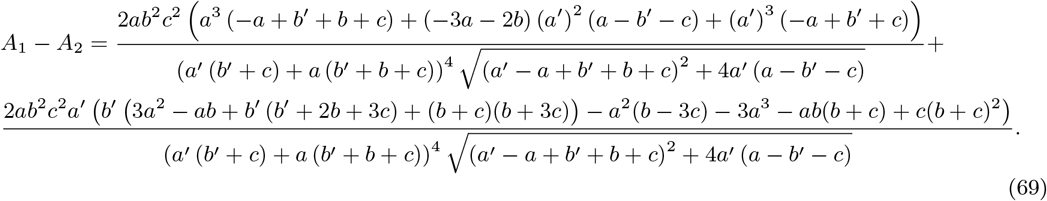

Note that *A*_1_ and *A*_2_ are the solution of the simultaneous equations Eqs. (68) and (69).

## D Derivation of the steady-state mean and variance of mature mRNA numbers for the mechanistic model

The statistics of mature mRNA numbers can be derived much more straightforwardly than those of the active number of Pol II. In steady-state, the flux across a system of species connected by irreversible reactions will be the same for each species and hence deletion of an intermediate species has no effect on the statistics of a downstream species. Hence, for the purpose of studying mature mRNA statistics in the steady-state [36], instead of the full scheme (1), we can consider a reduced scheme where the active Pol II is not explicitly described:

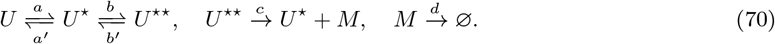

The stoichiometric matrix and the propensity vector are given by:

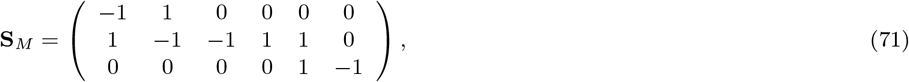

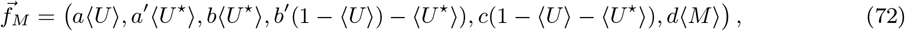

where 〈*X*〉 denotes the average number of molecules of species *X*. The species are numbered in the order *U, U ^⋆^, M* and the reactions in the order *U* → *U*^⋆^, *U*^⋆^ → *U, U*^⋆^ → *U*^⋆⋆^, *U*^⋆⋆^ → *U*^⋆^, *U*^⋆⋆^ → *U*^⋆^ + *M, M* → ⌀. We have here also used the same conservation law as in the previous Appendix C.

The time-evolution equations for the mean numbers and covariance matrix are given by Eqs. (46) and (47) where we replace **S** by **S**_*M*_ and 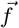 by 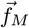. Setting the time derivatives to zero and solving these equations simultaneously, we find the steady-state mean and variance of mature mRNA given by 〈*n*_3_〉 and *C*_33_, respectively. The Fano factor of mature mRNA is then determined by their ratio:

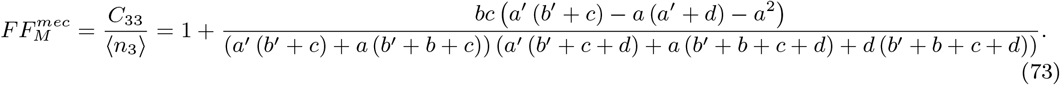

## E Comparison to reduction methods using number statistics

Here we compare the waiting time moment matching approach to two well-known model reduction techniques, which are: (i) matching of the moments of the number distributions and (ii) matching of the number distributions.

The first method consists of matching the first three moments of transcript number distributions of the mechanistic and two-state models. We find the steady-state mean, variance and skewness of the transcript numbers in two-state models as functions of *ρ*, *σ_b_* and *σ_u_* and then we equate them to the steady-state mean, variance and skewness of the transcript numbers computed for the mechanistic model (the first two moments are in Appendices C and D while the third moments can be computed similarly by solving the moment equations). For a given set of parameters of the mechanistic model, we solve the resulting system of three equations (with three unknowns *ρ*, *σ_b_* and *σ_u_*) numerically – this gives us the effective parameters of the two-state models. We have searched for numerical solutions throughout huge ranges of parameter space, however as shown in Fig. 7(A-C) and (G-I) we only find a physically meaningful solution (positive real numbers for the parameters of the two-state models which are shown by black dots in the figure) in the region of space given by Eq. (6) (where the mechanistic and two-state models can be matched using waiting time statistics; this is the region above the black solid line in the figure). Additionally note that the moment expressions for active Pol II for both the delayed telegraph model and mechanistic model are complicated and moment matching results in transcendental equations for the parameters *ρ*, *σ_b_* and *σ_u_*. The effective parameters for the delayed telegraph model, close to the contour lines where Δ = 0, are relatively small. Thus, conventional numerical solvers struggle to find solutions close to the boundaries (see Fig. 7(A-C)). In contrast, Fig. 7(G-I) show that effective parameters for the telegraph model can be found for nearly all mechanistic model parameter sets within the region given by Eq. (6). This is thanks to the relative simplicity of the analytical expressions for the telegraph model (compared to the transcendental equations encountered for the delayed telegraph model). In both Fig. 7(A-C) and Fig. 7(G-I), we use the FindRoot function with the Newton-Raphson method in *Mathematica*, where we start very close to the parameter prediction of the waiting time moment matching method and do 2 × 10^4^ iterations with a working precision of 200. Starting with points far from the theoretical predictions, no physical solutions are possible.

The second method consists of matching the transcript number distributions of the mechanistic and two-state models via maximum likelihood estimation (MLE). The results are presented in Fig. 7(D-F) and Fig. 7(J-L). The likelihood function is given by

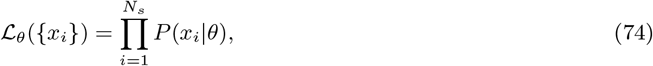

where {*x_i_*} is a set of samples (with length *N_s_*) generated using the delay SSA of the mechanistic model with specified parameters {*a, a*′, *b, b*′, *c*} (each *x_i_* represents the ith sample for the transcript number), *θ* is some candidate set of telegraph model parameters {*ρ, σ_u_, σ_b_*}, and *P* (*x_i_* | *θ*) is the probability of measuring *x_i_* given a telegraph model with parameters *θ*. We calculate the probability density function for a given parameter set *θ* using the exact solution for the delayed telegraph model [14] in Fig. 7(D-F) and the exact solution for the telegraph model [1] in Fig. 7(J-L). Next, we minimise the negative log-likelihood function

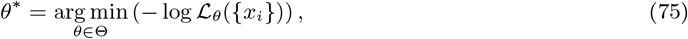

using the adaptive differential evolution algorithm in *Julia*’s `BlackBoxOptim` package to find the optimal parameters *θ*\* of the two-state model, where Θ is the set of all possible two-state model parameters (essentially amounting to a choice of parameter space bounds in `BlackBoxOptim`). Using these optimal parameters we obtain the steady-state distributions of active Pol II and of mature mRNA using the exact solutions of the two-state models. Finally, we compute the Hellinger distance (*h*) between these distributions and the corresponding ones from the mechanistic model – the distance is shown by the colour in Fig. 7(D-F) and Fig. 7(J-L). Note that in the regions where Δ > 0, *h* is generally smaller than in the regions where Δ < 0 i.e., the transcript number distributions found by MLE best approximate those of the mechanistic model in the region of parameter space given by Eq. (6) (where the mechanistic and two-state models can be matched using waiting time statistics). Therefore, we can conclude that waiting time moment matching agrees with this alternative model reduction method, albeit *the former is much more computationally efficient and accurate than the latter*.

